# Stress-Induced Iron-Sulfur Cluster Damage as a Conserved Trigger of the Stringent Response

**DOI:** 10.1101/2025.07.29.666095

**Authors:** Eva Michaud, Lorena Ricci, Claire Lallement, Lars Barquist, Vincent Cattoir, Charlotte Michaux, Régis Hallez, Séverin Ronneau

## Abstract

Pathogenic bacteria rely on the stringent response to adapt to the complex and fluctuating conditions encountered within the host. However, the mechanisms by which the stringent response senses host-induced stress remain poorly understood. Here, we identify iron–sulfur (Fe–S) cluster damage as a conserved trigger of the stringent response in major Gram-negative pathogens, including *Salmonella enterica*, *Enterobacter cloacae*, and *Klebsiella pneumoniae*. We demonstrate that Fe–S cluster disruption—caused by oxidative stress or metal imbalance—restricts the intracellular pools of sulfur-containing and branched-chain amino acids, thereby activating the ribosome-associated (p)ppGpp synthetase RelA. Furthermore, we show that iron availability governs recovery from Fe–S cluster damage, modulating the dynamics of the stringent response. Finally, we emphasize the dual role of (p)ppGpp in transcriptional regulation, enhancing bacterial fitness during Fe–S cluster stress while simultaneously promoting virulence by upregulating the SPI-2 type III secretion system. Together, these findings uncover a conserved mechanism by which pathogenic bacteria integrate metabolic stress into adaptive gene regulation and virulence, positioning Fe–S cluster integrity as a pivotal node linking environmental sensing to transcriptional control during infection.

## Introduction

From the first contact with their host, pathogenic bacteria encounter stress. During infection, these pathogens face numerous challenges, including exposure to reactive oxygen and nitrogen species (ROS and RNS, respectively), osmotic pressure, and fluctuations in pH and oxygen levels (1). Additionally, hosts employ nutritional immunity to restrict bacterial growth by manipulating metal availability. For instance, host cells can sequester essential metals such as iron by secreting chelating agents or actively removing them via specific transporters. Beyond metal sequestration, hosts can also intoxicate invading bacteria with toxic metal concentrations as an antimicrobial response (2–5).

To thrive in such challenging environments, bacterial pathogens have evolved sophisticated strategies to sense and adapt to host-induced stresses (1). One of the most universal mechanisms is the stringent response, which is mediated by the alarmone guanosine tetra- or penta-phosphate, collectively known as (p)ppGpp. This second messenger transiently alters bacterial physiology by regulating critical processes such as transcription, translation, and DNA replication (6–10). By tightly controlling (p)ppGpp levels, bacteria can balance growth and survival, making the stringent response a pivotal component of bacterial adaptation (11, 12). Consistently, mutants that fail to properly regulate (p)ppGpp struggle to proliferate in host tissues and exhibit poor survival during infection (7).

The intracellular levels of (p)ppGpp are primarily controlled by a conserved family of proteins known as RelA-SpoT Homologs (RSH) (11, 12). These proteins typically feature a multi-domain architecture, with two N-terminal enzymatic domains responsible for (p)ppGpp synthesis and degradation, and a C-terminal regulatory tail that integrates environmental signals. Most bacteria regulate (p)ppGpp levels using a single bifunctional RSH protein, but some species harbor multiple copies. In γ-proteobacteria, including *Salmonella enterica* serovar Typhimurium (henceforth *Salmonella*), (p)ppGpp levels are controlled by two proteins: RelA, a dedicated (p)ppGpp synthetase, and SpoT, which has both synthetase and hydrolase activities. RelA primarily senses amino acid availability during translation. Under amino acid starvation, RelA associates with uncharged tRNAs at the ribosomal A-site, triggering (p)ppGpp synthesis (13). SpoT, on the other hand, integrates diverse environmental signals, including fatty acid starvation and carbon source downshift, to modulate (p)ppGpp levels (14–16). A large body of evidence suggests a link between (p)ppGpp metabolism and metal availability across diverse bacterial species, including *Escherichia coli* (17), *Salmonella enterica* (18), *Bacillus subtilis* (19), *Enterococcus faecalis* (20), *Streptococcus pneumoniae* (21), and *Staphylococcus aureus* (22). However, the molecular mechanisms by which metal availability regulates the intracellular pool of (p)ppGpp remain poorly understood.

Here, by investigating how metal availability influences (p)ppGpp levels, we reveal that disruption of iron–sulfur (Fe–S) clusters activates the stringent response in several pathogenic γ-proteobacteria, including *Salmonella enterica*, *Enterobacter cloacae*, and *Klebsiella pneumoniae*. We demonstrate that Fe–S cluster damage leads to the depletion of branched-chain (isoleucine, leucine, and valine; ILV) and sulfur-containing (cysteine and methionine; CM) amino acids, thereby enhancing the activity of the ribosome-associated (p)ppGpp synthetase RelA. Furthermore, we show that diverse environmental stressors—such as fluctuations in metal availability and oxidative stress—modulate the stringent response by compromising Fe–S cluster homeostasis. Finally, we establish that (p)ppGpp promotes bacterial fitness under Fe–S cluster stress and reprograms transcriptional networks to adapt to these hostile conditions. Together, our findings define a mechanistic link between Fe–S cluster damage and (p)ppGpp signaling, revealing how bacterial pathogens dynamically rewire their physiology in response to environmental challenges.

## Results

### Manganese stress triggers (p)ppGpp accumulation by restricting the pool of amino acids

Since the stringent response has been associated with metal homeostasis in numerous bacterial species (17–22), we hypothesized that fluctuations in metal availability could trigger (p)ppGpp accumulation. To test this, we first identified conditions where *Salmonella* experiences metal stress, either through starvation or excess, using growth as an indicator of metal-induced stress. In the absence of stress, the wild-type (WT) strain reaches stationary phase approximately 6 hours (T6) after dilution into fresh MOPS minimal medium (Fig. S1A). We therefore used T6 as a reference time point to evaluate the induction of metal stress under various conditions (Fig. S1B).

To induce metal starvation, we first prepared media lacking magnesium (-Mg), iron (-Fe), zinc (-Zn), copper (-Cu), manganese (-Mn), or cobalt (-Co). Surprisingly, only Mg depletion significantly reduced bacterial growth at T6 (Fig. S1B), suggesting that residual trace metals were sufficient to support *Salmonella* growth in the other conditions. In support of that, the presence of DPI, TPEN, and BCS chelators (23–25) impaired bacterial growth, suggesting successful starvation for Fe, Zn, and Cu, respectively. To confirm the specificity of each chelator, we verified that the addition of the corresponding metal was sufficient to restore bacterial growth in the presence of the cognate chelator (Fig. S1B). For Mn and Co, we were unable to identify any specific chelators for which the exogenous addition of the corresponding metal could restore growth. Beyond starvation, we also assessed whether metal excess (+Mg, +Fe, +Zn, +Cu, +Mn, +Co) was toxic to *Salmonella*. We found that excess Zn, Cu, Mn and Co severely inhibited bacterial growth, while supplementation with elevated levels of Mg or Fe (1 mM) did not affect growth under our experimental conditions (Fig. S1C).

To investigate whether metal stress influences the stringent response, we monitored (p)ppGpp accumulation using thin-layer chromatography (TLC) under those metal stress conditions. To prevent indirect effects of stress on (p)ppGpp accumulation due to reduced uptake of radioactive ³²P, we first cultivated *Salmonella* with ³²P under non-stress conditions (MOPS). This approach ensured efficient incorporation of the radiolabel into the nucleotide pool, as evidenced by successful labeling of GTP (first lane in Fig. 1A). Then, we diluted the radiolabeled *Salmonella* into stress-inducing conditions and monitored variations in (p)ppGpp levels. Of the eight conditions tested, only the excess of Mn triggered (p)ppGpp accumulation, independently of its associated counterion (Fig. 1A, Fig. S1D).

**Fig. 1.**
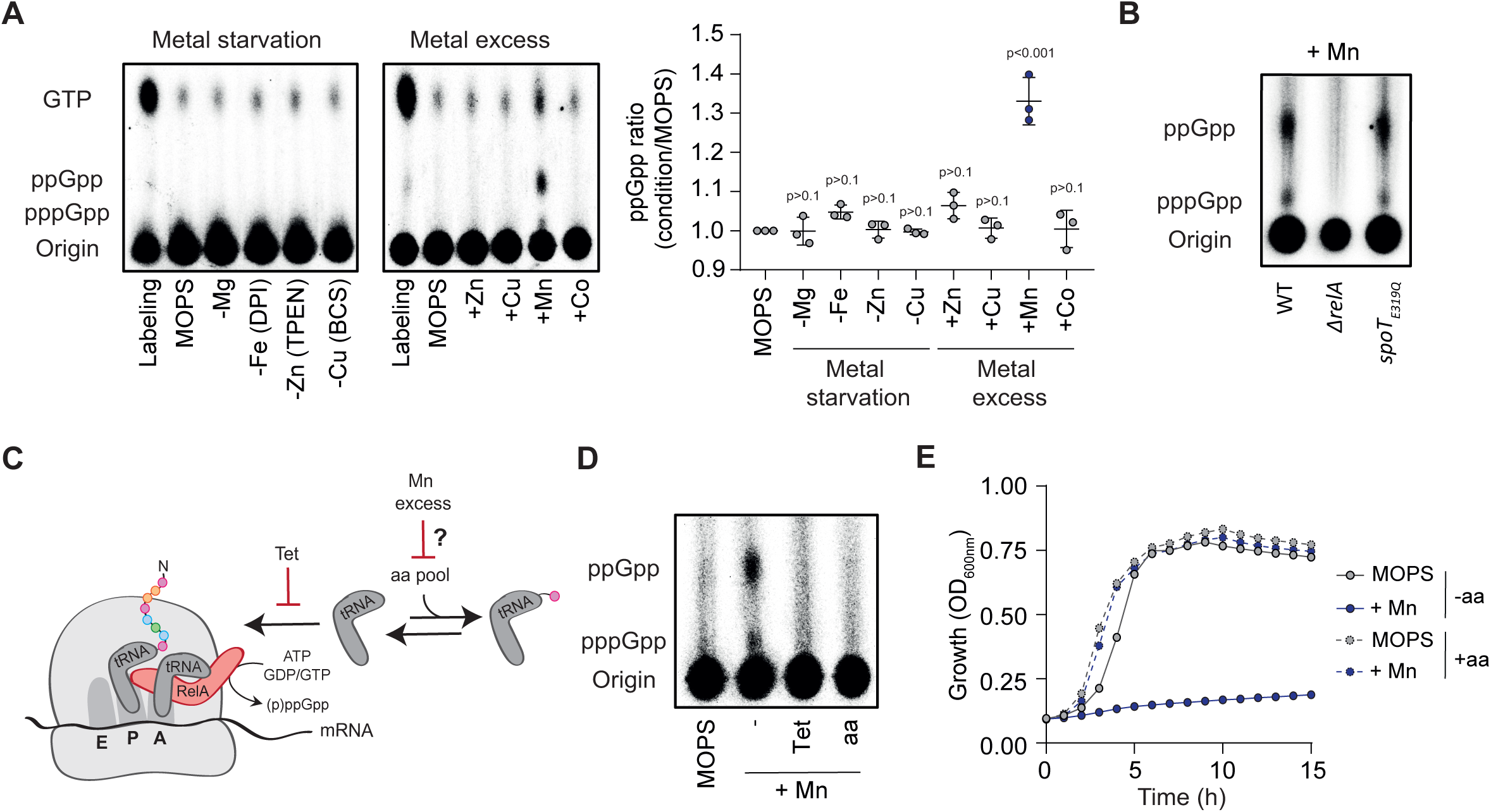
Manganese stress triggers (p)ppGpp accumulation through RelA. **(A)** Representative thin-layer chromatography (TLC) autoradiogram (left) and quantification (right) of ³²P-labeled nucleotide extracts from wild-type (WT) *Salmonella*. The first lane corresponds to labeled cells prior to dilution into MOPS minimal medium (unstressed) or metal stress conditions. **(B)** Representative TLC autoradiogram of nucleotide content in WT, Δ*relA*, and *spoT*_E319Q_ strains incubated with 1 mM MnSO₄ (+Mn) for 30 min. **(C)** Proposed model of RelA activation in response to amino acid limitation. **(D)** Representative TLC autoradiogram of nucleotide content in WT cells incubated for 30 min in MOPS (unstressed) or in the presence of 1 mM MnSO₄ (+Mn), with or without additional treatment: no supplementation (−), tetracycline (Tet), or a mixture of 20 amino acids (aa). **(E)** Growth curves of WT cells in MOPS minimal medium with or without 1 mM MnSO₄ (+Mn) and/or 20 amino acids (aa). *p* values are indicated (ANOVA with Dunnett’s correction in A); error bars represent mean ± SD; *n* ≥ 3. Each autoradiogram shown is representative of at least three independent experiments.

To determine whether (p)ppGpp was produced by RelA or SpoT, we monitored (p)ppGpp levels in a *relA* mutant (Δ*relA*) and in a strain containing a synthetase-dead version of SpoT (*spoT_E319Q_*) in the presence of elevated levels of Mn (26). Strikingly, the absence of RelA completely abolished (p)ppGpp accumulation during Mn intoxication, whereas no difference was observed between the WT and the *spoT_E319Q_* mutant (Fig. 1B). These results suggest that Mn stress leads to the activation of RelA synthetase activity likely by triggering accumulation of uncharged tRNA (Fig. 1C). In agreement with this hypothesis, the addition of tetracycline, an antibiotic that prevents RelA activation by blocking the entry of uncharged tRNAs into the ribosomal A-site (27, 28), completely abolished (p)ppGpp accumulation during Mn excess (Fig. 1C-1D). Furthermore, supplementation with amino acids completely abolished (p)ppGpp accumulation and rescued the associated growth defect of Mn-intoxicated cells (Fig. 1D-E). Together, these results demonstrate that excess Mn triggers RelA-dependent (p)ppGpp production by restricting the pool of amino acids.

### Disruption of Fe-S clusters leads to (p)ppGpp accumulation in Mn-intoxicated cells

Previous studies have suggested that Mn toxicity results from its ability to outcompete Fe for proteins that cannot use Mn as a cofactor (29, 30). For instance, Mn excess has been shown to interfere with heme synthesis, leading to depletion of cytochrome oxidase activity and ultimately to the failure of aerobic respiration (29). To determine whether Mn intoxication can trigger the stringent response by impairing bacterial respiration, we measured (p)ppGpp levels in the presence of potassium cyanide (KCN). KCN inhibits the cytochrome oxidase by binding with high affinity to the iron atom in its catalytic center, thereby blocking electron transfer to oxygen (31). As previously observed for Mn excess, inhibition with KCN triggers the accumulation of (p)ppGpp through RelA (Fig. S2A). Moreover, this accumulation was completely abolished by the addition of tetracycline or amino acids, as shown for Mn excess (Fig. S2B). If the inhibition of cytochrome oxidase activity is responsible for the accumulation of (p)ppGpp, we reasoned that a strain lacking all three terminal cytochrome oxidases (Δ*cyoA*Δ*cydB*Δ*appB*) should grow poorly and constitutively display high levels of (p)ppGpp. Although this strain exhibits a severe growth defect (Fig. S2C), we did not observe any increase in (p)ppGpp levels in the absence of stress (Fig. S2D). Moreover, this strain was still able to accumulate (p)ppGpp in the presence of KCN or excess Mn (Fig. S2D), demonstrating that inhibition of cytochrome oxidases cannot explain (p)ppGpp accumulation upon KCN or Mn intoxication.

Beyond heme synthesis, we reasoned that Mn might also directly interfere with Fe-S cluster-containing proteins. In agreement with this hypothesis, it has been shown that disruption of Fe-S clusters causes multiple auxotrophies (32), a phenotype we also observed under Mn intoxication (Fig. 1E). This effect can be explained, at least in part, by the inactivation of the Fe-S-dependent enzymes dihydroxyacid dehydratase (IlvD) and isopropylmalate dehydratase (LeuC), which are involved in the biosynthesis of the branched-chain amino acids isoleucine, leucine, and valine (ILV) (28, 33–35) (Fig. 2A). We therefore tested whether ILV supplementation could restore growth under Mn stress. Supplementation with branched amino acids poorly rescued growth during Mn stress, supporting that inactivation of other Fe-S cluster-containing proteins contributes to Mn toxicity (Fig. 2B). Since Fe-S cluster biogenesis requires sulfur atoms from amino acids (Fig. 2A), we wondered whether the addition of the sulfur-containing amino acids cysteine (C) and methionine (M) could reduce Mn toxicity (Fig. 1E). In contrast with ILV supplementation, we observed that addition of CM substantially rescued growth in the presence of elevated Mn levels (Fig. 2B), suggesting that sulfur-containing amino acids mitigate Mn toxicity by promoting the regeneration of a functional pool of Fe-S clusters. To test this hypothesis, we deleted *hscA*, which encodes a chaperone involved in Fe-S cluster maturation (36), and tested whether CM supplementation could still rescue bacterial growth under Mn excess (Fig. 2A). Strikingly, the growth defect of Δ*hscA* under Mn excess was not rescued by supplementation with sulfur-containing amino acids (Fig. 2C), supporting that cysteine and methionine mitigate Mn toxicity by supplying sulfur atoms required for Fe-S cluster assembly. In addition, the *hscA* mutant also displayed a higher susceptibility than the WT to Mn toxicity (Fig. S3A-B), which emphazise the importance of Fe-S assembly during Mn excess. Remarkably, the combined supplementation of both branched-chain (ILV) and sulfur-containing (CM) amino acids provided the most effective protection against Mn-induced growth inhibition (Fig. 2B). These findings suggest that Mn-induced damage to Fe-S clusters results in multiple auxotrophies: one arising from the loss of Fe-S-dependent enzymes required for ILV biosynthesis, and another reflecting increased cellular demand for sulfur donors to support Fe-S cluster synthesis.

**Fig. 2.**
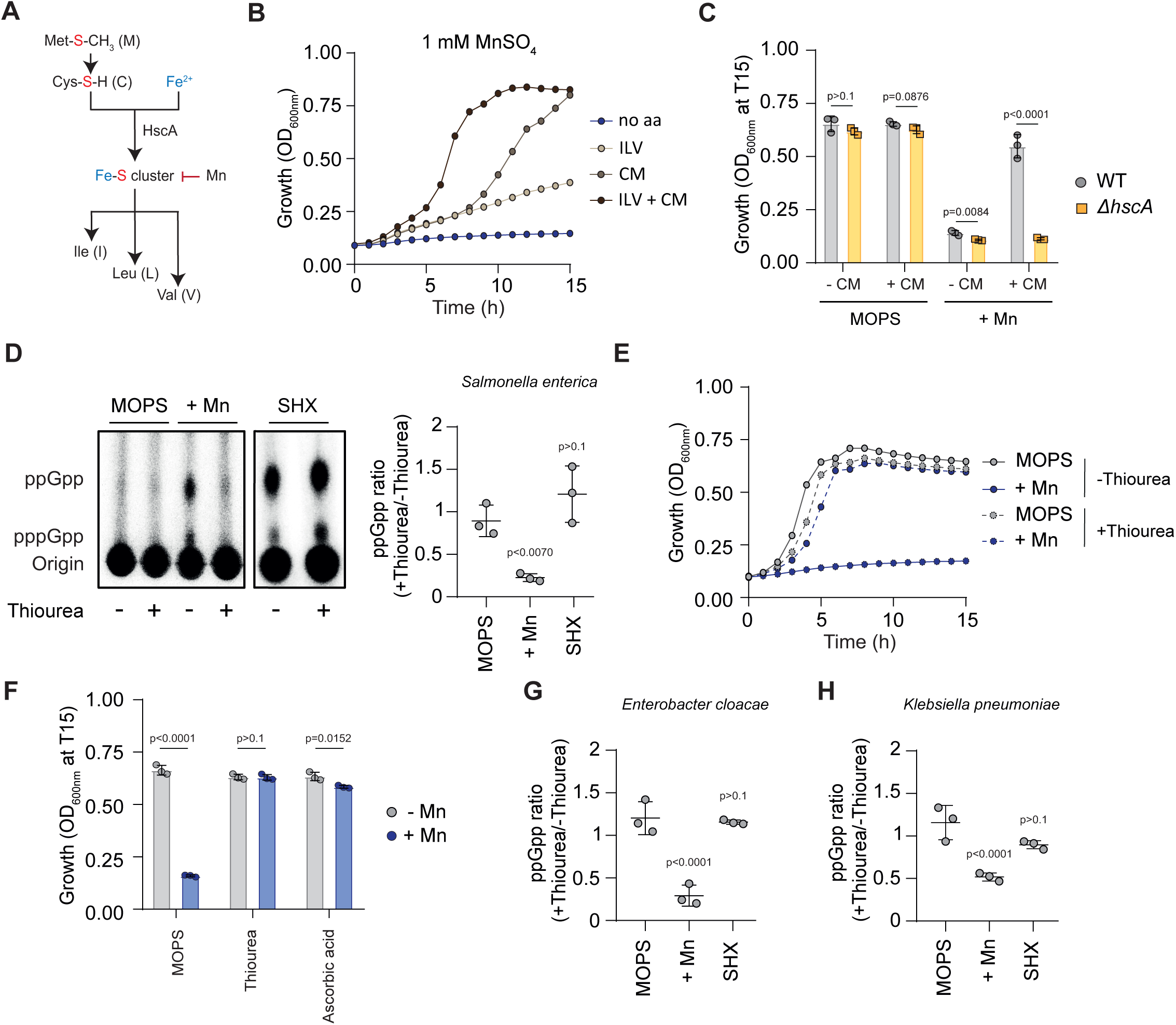
Manganese intoxication leads to Fe-S cluster damage. **(A)** Proposed model of Fe-S cluster assembly and its connection to amino acid metabolism. **(B)** Growth curves of WT cells in MOPS minimal medium containing 1 mM MnSO₄ and with or without amino acid (aa) supplementation. ILV: isoleucine, leucine, valine; CM: cysteine and methionine. **(C)** Optical density (OD₆₀₀_nm_) of Δ*hscA* strains after 15 h (T15) of growth in MOPS minimal medium in presence or absence of 1 mM MnSO₄ and with (+) or without (-) cysteine and methionine (CM) supplementation. **(D)** Representative TLC autoradiogram (left) and quantification (right) of ³²P-labeled nucleotide extracts from WT *Salmonella* grown for 30 min in MOPS minimal medium in presence or absence of 1 mM MnSO₄ (+Mn) or 0.4 mg/ml SHX and supplemented with (+) or without (-) 100 mM thiourea. **(E)** Growth curves of WT cells in MOPS minimal medium in presence or absence of 1 mM MnSO₄ and supplemented with or without 100 mM thiourea. **(F)** Optical density (OD₆₀₀_nm_) of WT *Salmonella* after 15 h (T15) of growth in MOPS minimal medium in presence or absence of 1 mM MnSO₄ and supplemented with or without 100 mM of thiourea or 50 mM ascorbic acid. **(G-H)** Quantification of relative ppGpp levels in WT *Enterobacter cloacae* (**G**) and WT *Klebsiella pneumoniae* (**H**) grown for 30 min in MOPS minimal medium in presence or absence of 1 mM MnSO₄ (+Mn) or 0.4 mg/ml SHX and supplemented with or without 100 mM thiourea. *p* values are indicated (multiple *t*-tests in C and F, ANOVA with Dunnett’s correction in D, G and H); error bars represent mean ± SD; *n* = 3. Each autoradiogram shown is representative of at least three independent experiments.

Since reducing environments promote Fe-S cluster assembly and stability *in vitro* (37, 38), we hypothesized that the addition of thiourea—a commonly used reducing agent—might help bacteria preserve a functional pool of Fe-S cluster-containing proteins, thereby limiting the subsequent activation of RelA. Consistent with this hypothesis, we observed that thiourea abolished (p)ppGpp accumulation during Mn stress (Fig. 2D). Interestingly, thiourea also prevented the elevation of (p)ppGpp levels in response to KCN (Fig. S3C), which also interferes with Fe-S cluster-containing proteins due to its strong binding to iron atoms (39). As a control, we confirmed that thiourea had no effect on (p)ppGpp accumulation induced by serine hydroxamate (SHX), a serine analog that triggers amino acid starvation by inhibiting seryl-tRNA synthetase (40) (Fig. 2D). In line with these assays, thiourea also alleviated the growth defects associated with excess Mn, supporting the functionality of Fe-S cluster-containing proteins under these conditions (Fig. 2E). Likewise, another reducing agent — ascorbic acid — also effectively suppressed Mn-induced growth defects (Fig. 2F). Thus, these data support the role of Fe-S cluster status in regulating (p)ppGpp levels in *Salmonella*.

Next, we investigated if other clinically relevant Gram-negative pathogens could modulate the stringent response in response to Fe-S cluster damage. To test this, we measured (p)ppGpp levels in two other pathogenic γ-proteobacteria, *Enterobacter cloacae* and *Klebsiella pneumoniae*, in the presence or absence of Mn excess. Similar to our findings in *Salmonella*, we observed that Mn excess and SHX induced (p)ppGpp accumulation in these species (Fig. 2G-H, S3D-E). Moreover, supplementation with thiourea completely abolished (p)ppGpp accumulation during Mn stress, but not in the presence of SHX (Fig. 2G-H, S3D-E). Collectively, these results indicate that the regulation of the stringent response by Fe-S cluster damage is not limited to *Salmonella* but is also conserved across other clinically relevant pathogens.

### Iron limitation promotes the stringent response upon conditions inducing Fe-S cluster damage

Having identified that the disruption of Fe-S clusters homeostasis can elevate (p)ppGpp levels, we wondered why iron starvation did not lead to (p)ppGpp accumulation under our experimental conditions (Fig. 1A). One hypothesis is that preexisting Fe-S clusters are sufficient to maintain amino acid metabolism during a brief iron starvation, and that the destabilization of these preexisting clusters is required to trigger the stringent response. Since iron is a critical component of Fe-S clusters (Fig. 2A), we hypothesized that limiting iron availability might still regulate the stringent response, but only in the presence of stress that destabilizes Fe-S clusters and requires the assembly of new ones.

To investigate this hypothesis, we first identified a condition that induces a transient disruption of Fe-S clusters which allows bacteria to resume growth after stress. We found that the addition of 0.5 mM Mn significantly extends the lag phase of *Salmonella* but allows recovery, supporting that bacteria successfully restored a functional pool of Fe-S clusters (Fig. 3A). Under these conditions, (p)ppGpp levels rise rapidly in response to Mn excess and return to their pre-stress levels within two hours (Fig. 3B). In contrast, higher concentrations of Mn (1 mM) were bacteriostatic and led to the maintenance of elevated levels of (p)ppGpp over time (Fig. 3B). Next, we determined how iron availability might affect the recovery of *Salmonella* after mild Mn excess. To reduce iron availability in the medium, we used a concentration of the Fe-chelator DPI that does not affect bacterial growth in the absence of stress (Fig. 3C). Strikingly, limiting iron availability using DPI drastically increased the lag phase following mild Mn excess and promoted the maintenance of the stringent response, as observed at higher concentrations of Mn (Fig. 3C-D).

**Fig. 3.**
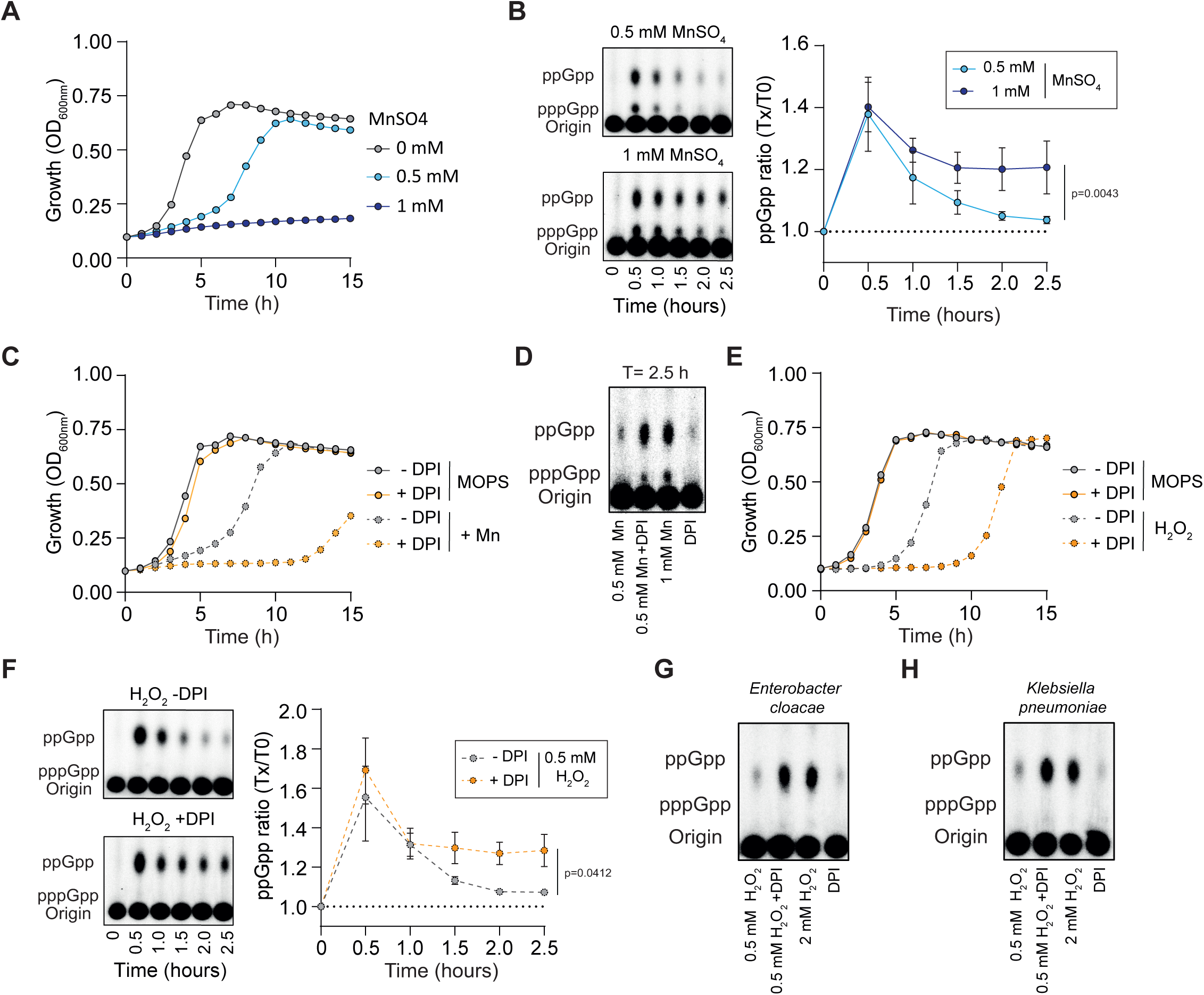
Iron availability governs the stringent response under Fe–S cluster-damaging stress. **(A)** Growth curves of WT *Salmonella* in MOPS minimal medium supplemented with 0, 0.5, or 1 mM MnSO₄. **(B)** Representative TLC autoradiograms (left) and quantification (right) of ³²P-labeled nucleotide extracts from WT *Salmonella* over time following addition of 0.5 or 1 mM MnSO₄. **(C)** Growth curves of WT cells in MOPS minimal medium with 0 or 0.5 mM MnSO₄, and with or without 0.1 mM DPI. **(D)** Representative TLC autoradiogram of ³²P-labeled nucleotide extracts from WT cells grown for 2.5 hours in MOPS minimal medium with 0.5 or 1 mM MnSO₄ and 0 or 1 mM DPI. **(E)** Growth curves of WT cells in MOPS minimal medium with 0 or 0.5 mM H₂O₂, in the presence or absence of 0.1 mM DPI. **(F)** Representative TLC autoradiograms (left) and quantification (right) of ³²P-labeled nucleotide extracts from WT *Salmonella* over time after treatment with 0.5 mM H₂O₂, with or without 1 mM DPI. **(G–H)** Representative TLC autoradiograms of ³²P-labeled nucleotide extracts from WT *E. cloacae* (**G**) and *K. pneumoniae* (**H**) grown in MOPS minimal medium with 0.5 or 2 mM H₂O₂, and supplemented with 0 or 1 mM DPI for 2.5 h. *p* values in D and F were determined by unpaired *t*-test (final time point); error bars represent mean ± SD; *n* = 3. Each autoradiogram shown is representative of at least three independent experiments.

We next wondered whether iron limitation might also extend the stringent response under infection-relevant stresses that destabilize Fe-S clusters. To explore this, we examined the role of iron availability during oxidative stress, which destabilizes Fe-S clusters (41, 42) and elevates ppGpp levels via RelA by generating an amino acid starvation (43) (Fig. S4A-C). In line with our model, iron limitation drastically delayed bacterial regrowth following mild H₂O₂ stress and led to the maintenance of elevated levels of ppGpp over time (Fig. 3E-F). Remarkably, both phenotypes were also observed during more severe H₂O₂ stress (Fig. S4D-E). Given the similar response of *Salmonella* treated with Mn excess or H₂O₂, we examined whether Mn excess induces oxidative stress under our experimental conditions. Using the P*_katG__gfpOVA* ROS-specific biosensor (44) in the WT strain, we observed robust GFP accumulation following H₂O₂ treatment but no increase after Mn exposure (Fig. S4F), indicating that Mn stress does not generate oxidative stress in our experimental conditions.

To extend our findings to other pathogens, we performed the same approach for *Enterobacter cloacae* and *Klebsiella pneumoniae*, both of which accumulating (p)ppGpp in response to Fe-S cluster damage (Fig. 2G-H, S3D-E). As shown for *Salmonella*, we observed that both species accumulate ppGpp – but not pppGpp – in response to H₂O₂ exposure and that limiting iron availability with DPI delayed bacterial recovery and promoted the stringent response, as observed when H₂O₂ concentration was higher (Fig. 3G-H, S5A-B). Collectively, these findings demonstrate that (p)ppGpp levels are regulated by iron availability, which dictates the ability of bacteria to regenerate a functional pool of Fe–S clusters following stress exposure.

### Elevated (p)ppGpp levels reprogram bacterial transcription in response to Fe-S cluster damage

Our previous data demonstrated that the Fe-S cluster damage dynamically modulates (p)ppGpp levels. However, whether activation of the stringent response contributes to bacterial fitness under these conditions remained unclear. To address this, we compared the growth of WT and Δ*relA* mutant strains under conditions known to induce Fe-S cluster damage, such as Mn excess or oxidative stress. The Δ*relA* mutant failed to grow under stress conditions at concentrations that still permitted growth of the WT strain. (Fig. 4A). We also tested the ability of both strains to produce an inducible GFP in the presence or absence of stress. While GFP production was comparable between strains without stress, both Mn excess and H₂O₂ completely abolished GFP synthesis in the Δ*relA* mutant but not in the WT strain, which sustained substantial GFP expression under stress (Fig. 4B). Together, these results demonstrate that (p)ppGpp accumulation enhances bacterial fitness following Fe-S cluster damage by maintaining essential cellular functions such as transcription and translation.

**Fig. 4.**
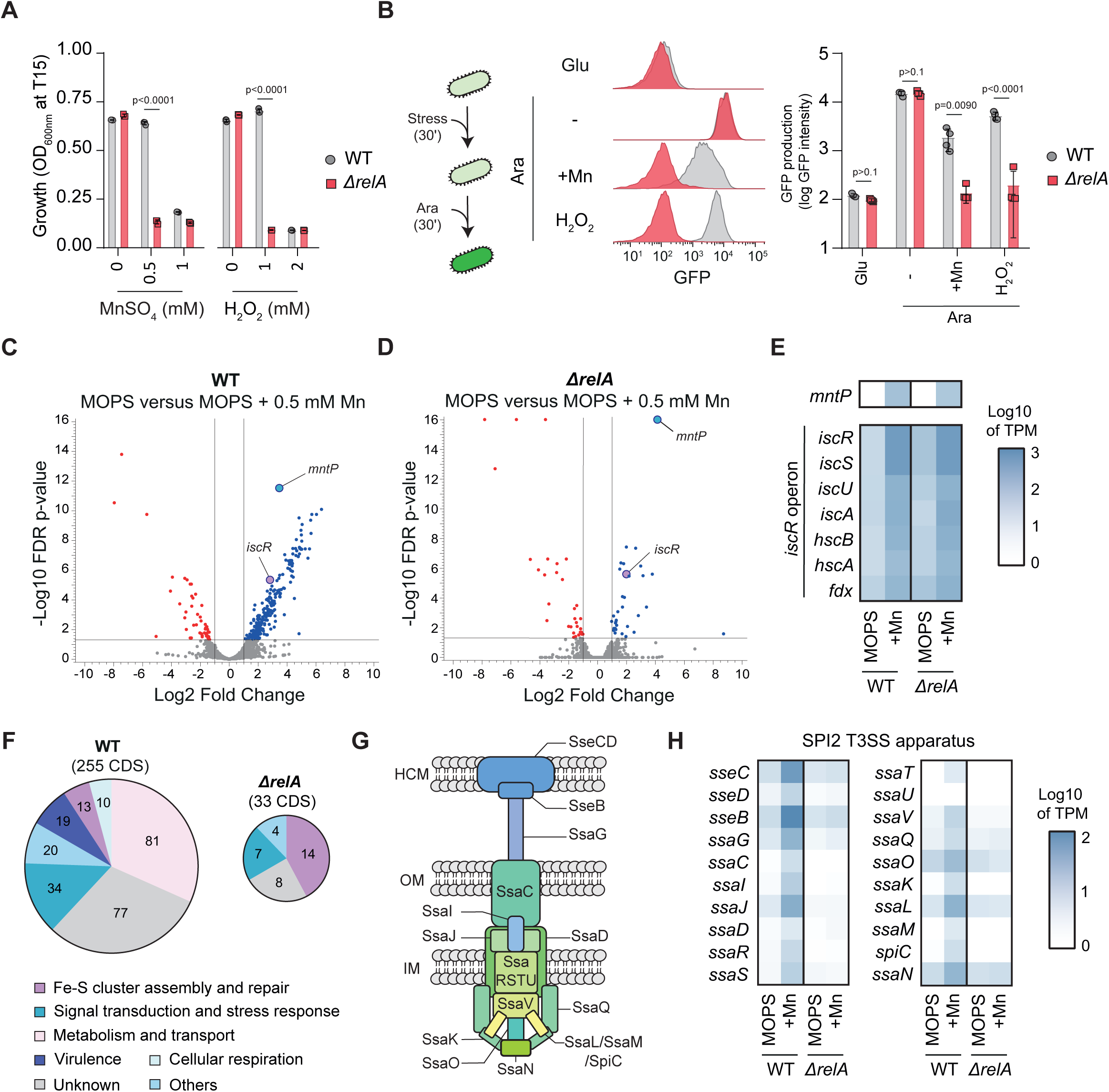
(p)ppGpp modulates stress tolerance and global gene expression during Fe-S cluster-damaging conditions. (A) Optical density (OD₆₀₀_nm_) of WT and *ΔrelA* strains after 15 hours (T15) of growth in MOPS minimal medium supplemented with increasing concentrations of MnSO₄ or H₂O₂. **(B)** Representative FACS plots and quantification of transcriptional and translational activity in WT and Δ*relA* strains grown in MOPS minimal medium with glucose (Glu) or arabinose (Ara), in the absence (-) or presence of 1 mM MnSO₄ (+Mn) or 0.5 mM H₂O₂. **(C-D)** Volcano plots showing differentially expressed genes (RNA-seq) in WT (**C**) and Δ*relA* (**D**) strains following 30 min exposure to 0.5 mM MnSO₄ compared to unstressed MOPS controls. **(E)** Heatmap showing expression of *mntP* and the *IscR* regulon in WT and *ΔrelA* strains in the absence (MOPS) or presence (+Mn) of 0.5 mM MnSO₄. **(F)** Pie charts showing functional categories of upregulated genes during Mn stress in WT and Δ*relA* strains. Numbers indicate in each category. **(G)** Simplified representation of the SPI2 T3SS apparatus. HCM, Host Cell Membrane; OM, Outer Membrane; IM, Inner Membrane. **(H)** Heatmap showing expression of the SPI2 T3SS apparatus genes in WT and *ΔrelA* strains in the absence (MOPS) or presence (+Mn) of 0.5 mM MnSO₄. *p* values were calculated using multiple *t*-tests (A-B); error bars represent mean ± SD; *n* ≥ 3.

Next, to elucidate how (p)ppGpp facilitates adaptation under Fe-S cluster damaging conditions, we performed RNA-seq on WT and Δ*relA* strains exposed to Mn excess. We selected Mn over H₂O₂ to avoid confounding effects from ROS-specific genes (*e.g., katG*) and used a mild Mn concentration (0.5 mM) that induces (p)ppGpp accumulation (Fig. 3B) while still permitting transcriptional activity in the *ΔrelA* mutant (Fig. S6A), thereby minimizing transcriptomic artifacts. Under these conditions, the WT strain significantly upregulated 255 coding sequences (CDS) and downregulated 55 CDS (Table S4), whereas the Δ*relA* mutant upregulated only 33 CDS and downregulated 32 CDS (Fig. 4C-E, S6B; Table S5). As expected, both strains similarly upregulated *mntP* (formely *yebN*), that encodes the main bacterial Mn efflux pump and which is under the control of the MntR regulator (47) (Fig. 4E). This confirmed that both strains experienced Mn stress and validated that our experimental conditions still allowed the Δ*relA* strain to mount a transcriptional response to environmental cues. Additionally, both strains upregulated the *iscR* regulon (Fig. 4E), which governs Fe-S cluster biogenesis (36) and is activated under conditions that compromise Fe-S cluster integrity (48). Together, these data ruled out the possibility that the increased Mn susceptibility of Δ*relA* results from defective expression of *mntP* or the *iscR* regulon.

Despite both strains upregulating *mntP* and the *iscR* regulon, the WT exhibited a broader transcriptional response (Fig. 4C-D, S6B; Table 4-5). Comparison with previous datasets (45) revealed that 64% of the CDS upregulated in the WT (162/255) belong to the RpoS regulon, a stress-related sigma factor positively regulated by (p)ppGpp (46) (Fig. S6C). Correspondingly, RpoS protein levels sharply increased during Fe-S cluster stress, including Mn excess or H₂O₂ exposure, but only in the presence of (p)ppGpp (Fig. S6D). These findings suggest that (p)ppGpp promotes activation of the RpoS regulon under Fe-S cluster damaging conditions.

Building upon the central role of (p)ppGpp in stress regulation, our transcriptomic analysis revealed significant upregulation of genes involved in substrate transport, metabolism, signal transduction, and stress responses (Fig. 4F). Importantly, we also observed elevated expression of the SPI-2 type III secretion system (SPI2 T3SS) apparatus (Fig. 4G-H), which is essential for intracellular survival within macrophages and the establishment of systemic infection (49, 50). This induction was exclusive to the WT strain and absent in the *relA* mutant (Fig. 4H), suggesting that perturbation of Fe-S cluster homeostasis may serve as a (p)ppGpp-mediated signal to promote SPI2 T3SS assembly during infection. Together, these findings highlight how (p)ppGpp enhances bacterial fitness by reprogramming transcription to support stress adaptation and virulence.

## Discussion

The stringent response is one of the most conserved adaptation mechanism, enabling bacterial pathogens to sense stress and reprogram gene expression to promote host colonization and survival (7). However, understanding how specific environmental cues encountered during infection modulate intracellular (p)ppGpp levels remains challenging although important given the therapeutic potential of disrupting this pathway to interfere with bacterial pathogenesis. Here, we show that Fe–S cluster damage is a key trigger of (p)ppGpp accumulation. Environmental stressors such as oxidative stress and metal imbalance compromise Fe–S cluster integrity, leading to specific amino acid auxotrophies, most notably for branched-chain (ILV) and sulfur-containing (CM) amino acids. These deficits result from inactivation of Fe–S-dependent enzymes, including IlvD and LeuC, and from increased sulfur demand required for Fe–S cluster repair. The ensuing metabolic perturbations promote RelA stabilization at the ribosomal A site, triggering a rapid burst of (p)ppGpp synthesis. This alarmone transiently reprograms bacterial transcription toward a stress-adaptive state, including induction of stress-responsive and virulence genes (Fig. 5). Importantly, this response is essential for maintaining core cellular processes, such as transcription and translation, under stress conditions. Moreover, we show that this mechanism is conserved across clinically relevant Enterobacteriaceae, including *Klebsiella pneumoniae* and *Enterobacter cloacae*.

**Fig. 5.**
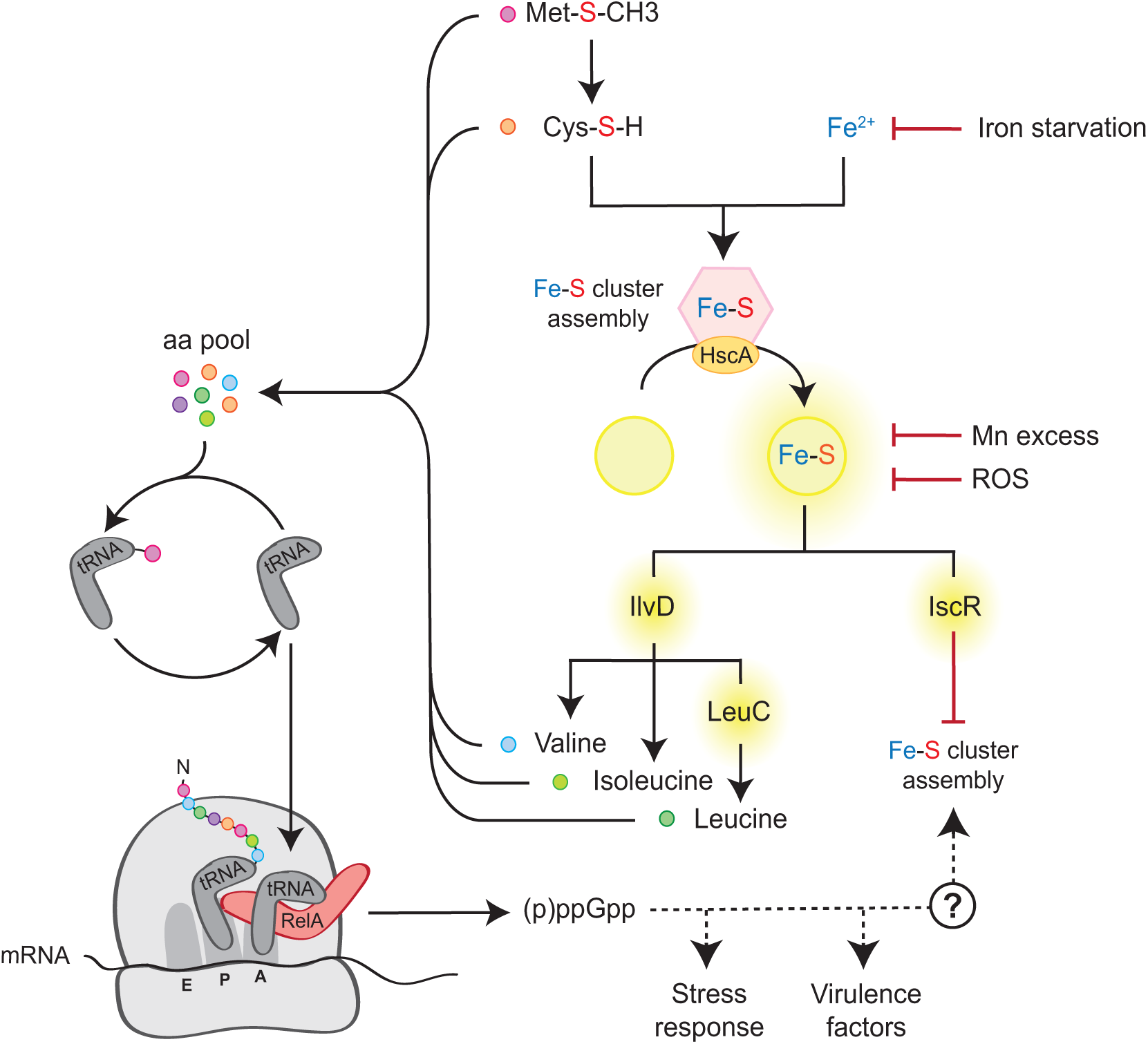
Model of (p)ppGpp regulation in response to Fe–S cluster damage. Environmental stressors such as oxidative stress and metal imbalance compromise Fe–S cluster integrity, resulting in amino acid auxotrophies—most notably for branched-chain (ILV) and sulfur-containing (CM) amino acids. These auxotrophies arise from the inactivation of Fe–S-dependent enzymes (*e.g.* IlvD, LeuC) and increased sulfur demand required for Fe–S cluster repair. The resulting amino acid limitation activates RelA, triggering a rapid accumulation of (p)ppGpp. This alarmone burst reprograms bacterial gene expression to promote stress adaptation and virulence.

By identifying Fe-S cluster damage as a trigger for (p)ppGpp induction, our study expands current understanding of how host-imposed stresses are converted into adaptive bacterial responses during infection. Pathogens are routinely exposed to Fe-S-damaging conditions, such as reactive oxygen and nitrogen species (ROS and RNS), metal fluctuations, and nutrient deprivation, especially within phagocytic immune cells (1). While the stringent response is classically associated with amino acid starvation, many pathogens are prototrophic and capable of synthesizing all amino acids. Our data support an expanded model in which (p)ppGpp functions as a broader stress integrator that responds not only to nutrient limitation but also to oxidative and nitrosative stress (28, 43). This broader view of (p)ppGpp signaling may explain why it has been implicated in oxidative stress resistance across a wide array of pathogens, including *Pseudomonas aeruginosa*, *Vibrio cholerae*, *Staphylococcus aureus*, and *Enterococcus faecalis* (20, 22, 51, 52). Moreover, we observed that (p)ppGpp accumulation following Fe-S cluster damage promotes the expression of *Salmonella* Pathogenicity Island 2 apparatus, which is essential for intracellular survival (49). These findings suggest that (p)ppGpp not only facilitates stress adaptation but also acts as a central hub that dynamically coordinates the expression of virulence determinants in response to environmental cues. The connection between Fe-S cluster integrity and stringent signaling highlights how pathogens exploit metabolic stress signals to fine-tune global regulatory networks during host colonization.

Mechanistically, we show that Fe-S cluster damage triggers RelA-dependent (p)ppGpp synthesis. Intriguingly, SpoT, the bifunctional (p)ppGpp synthetase/hydrolase, has been reported to respond to iron limitation in *Escherichia coli* (17), although the underlying mechanism remains elusive. This study raises the possibility of coordination between RelA and SpoT. A compelling hypothesis is that the hydrolase activity of SpoT is inhibited during Fe-S stress, allowing RelA-driven synthesis to proceed unopposed and prevent futile (p)ppGpp turnover. While direct evidence remains limited, other stress conditions suggest such interplay between (p)ppGpp enzymes. For example, during fatty acid starvation, RelA is activated via lysine depletion (53), whereas SpoT is regulated by its interaction with the acyl-carrier protein (ACP), which senses fatty acid intermediates (14, 15). Analogous regulatory crosstalk may occur under Fe–S stress, with RelA sensing Fe–S damage through amino acid limitation and SpoT directly sensing Fe availability to fine tune bacterial physiology, but this model requires further investigation.

The stringent response is associated with antibiotic tolerance, persistence, and biofilm formation, three major contributors to chronic infection and treatment failure (54–57). Elucidating how (p)ppGpp is regulated in response to metabolic stress opens new avenues to limit the success of bacterial infections. Specifically, targeting the molecular triggers or downstream effectors of (p)ppGpp signaling may sensitize pathogens to antibiotics or host immune clearance. This is particularly relevant in the context of slow-growing, metabolically altered bacterial populations that evade traditional antibiotics by entering a recalcitrant state (tolerance or persistence) during infection (58–60). Moreover, Fe-S cluster-damaging stresses that induce (p)ppGpp, such as ROS and RNS, have emerged as key determinants of antibiotic recalcitrance across numerous phylogenetically distant pathogens (61–66). However, the precise impact of (p)ppGpp on antibiotic recalcitrance under these conditions remains to be elucidated.

Our work provides a better understanding of how the stringent response is regulated by Fe-S cluster damage in bacterial pathogens. However, several open questions remain, such as how (p)ppGpp contributes to Fe-S cluster biogenesis or repair (Fig. 5). It may promote metabolic pathways that generate sulfur and iron precursors essential for Fe-S cluster assembly. However, we did not observe a significant (p)ppGpp-dependent regulation of ILV or CM biosynthetic genes, with the exception of *cdsH*, a cysteine desulfhydrase that could facilitate sulfur mobilization (Fig. S7A). Alternatively, (p)ppGpp may regulate iron homeostasis, as indicated by significant increased expression of iron-storage proteins such as FtnB and DPS in WT but not in the *relA* mutant (Fig. S7B). In constrast with Bfr and FtnA, the two other iron-storage protein of *Salmonella*, both FtnB and DPS have been linked to tolerance to oxidative stress and *Salmonella* virulence (67). It is noteworthy that (p)ppGpp also sharply increased the expression of the SPI-2 effector SpvB (Table S4), which may aid bacterial acquisition of host-derived iron (68). Lastly, approximately one-third of the (p)ppGpp-induced genes under Fe-S stress remain completely uncharacterized (Fig. 4E). Uncovering their roles could reveal new effectors of virulence, stress tolerance, or metabolic adaptation. As such, further investigation of (p)ppGpp-dependent transcriptional networks may not only deepen our understanding of bacterial stress adaptation but also identify promising targets for antimicrobial development.

Altogether, our study identifies Fe-S cluster damage as a previously underappreciated yet physiologically relevant trigger of the stringent response. This expands the canonical view of (p)ppGpp signaling beyond nutrient deprivation and highlights how intracellular pathogens exploit metabolic damage as a signal to activate stress responses and virulence programs. By revealing how environmental challenges are transduced into adaptive gene expression via the stringent response, our findings offer a new lens on bacterial pathogenesis and underscore (p)ppGpp metabolism as a promising target for therapeutic intervention.

## Acknowledgments

We thank all members of the Hallez lab, the BRM unit and Benjamin Ezraty for fruitful scientific discussions. We are also grateful to Jean-François Collet, Hélène Andrews-Polymenis, Michael McClelland, and Athanasios Typas for providing the *Salmonella* Single Gene Deletion library. We also thank Sophie Helaine for her critical reading of the manuscript.

This work was supported by the Marie Sklodowska Curie COFUND action (No 101034383) and the ATIP-Avenir program to S.R., the CARE ANR grant (22-PAMR-0002) to C.M., and the Welbio Starting Grant (WELBIO-CR-2019S-05) to R.H. R.H. is a senior research associates of the F.R.S. – FNRS.

## Authors contribution

S.R. conceived and designed the experiments. E.M., L.R., C.M., and S.R. performed the experiments. C.L., L.B. and C.M. analyzed the RNA-seq data. C.M., V.C. and R.H. provided intellectual input and contributed reagents, materials, and analysis tools. S.R. wrote the manuscript with input from all the authors.

## Declaration of interests

The authors declare no competing interests.

## Figure legends

**Fig. S1.**
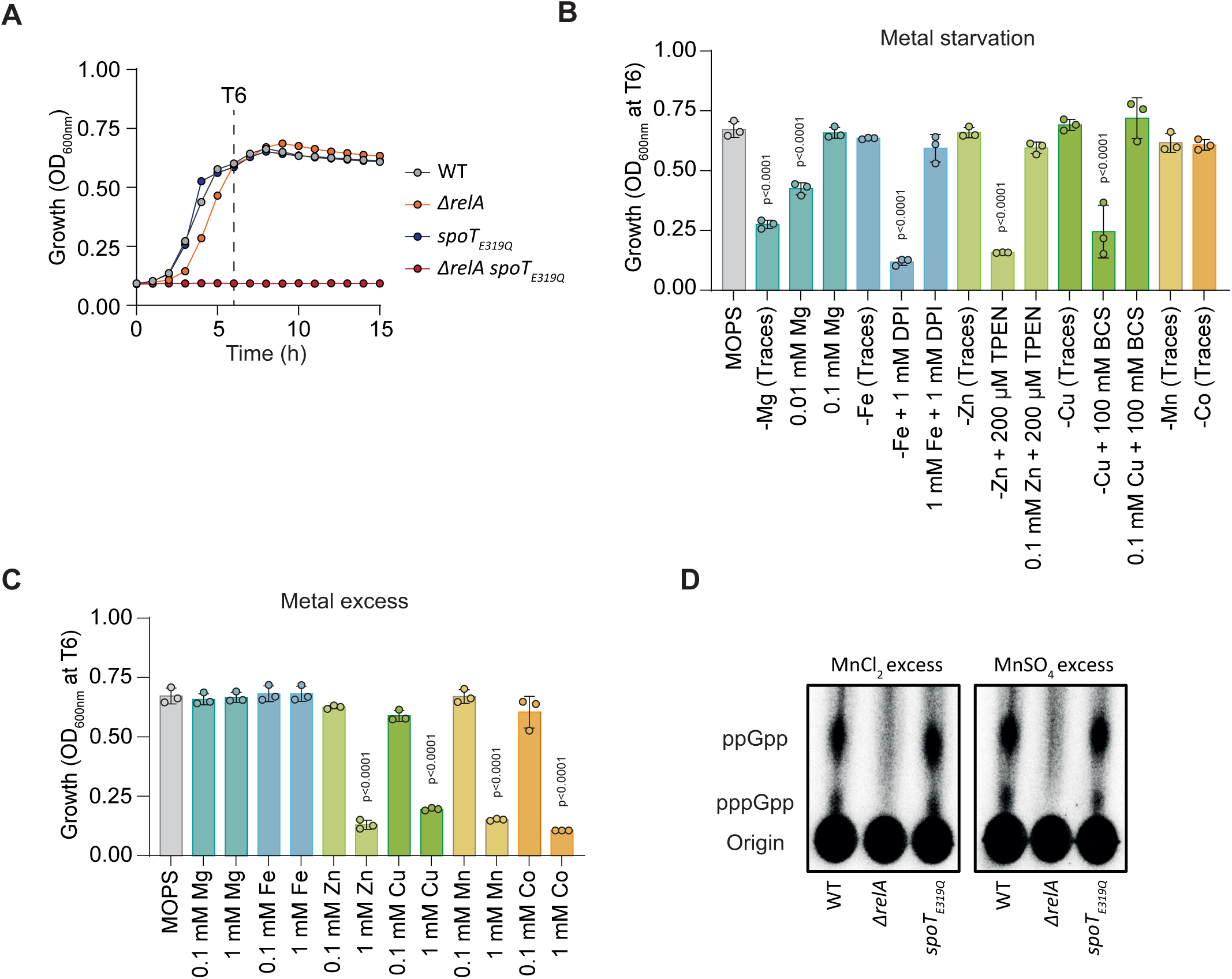
Interplay between (p)ppGpp metabolism and metal stress. **(A)** Growth curves of WT, *ΔrelA*, *spoT_E319Q_*, and *ΔrelA spoT_E319Q_ Salmonella* strains in MOPS minimal medium. **(B)** Optical density (OD₆₀₀_nm_) of WT *Salmonella* after 6 h (T6) of growth in MOPS minimal medium with or without magnesium (-Mg), iron (-Fe), copper (-Cu), zinc (-Zn), manganese (-Mn), or cobalt (-Co). Metal starvation was achieved using specific chelators: DPI (Fe), TPEN (Zn), and BCS (Cu). **(C)** Optical density (OD₆₀₀_nm_) of WT *Salmonella* after 6 h of growth in MOPS minimal medium supplemented with 0.1 or 1 mM of Mg, Fe, Zn, Cu, Mn, or Co. **(D)** Representative TLC autoradiogram showing nucleotide content in WT, *ΔrelA*, and *spoT_E319Q_* strains after treatment with 1 mM MnCl₂ or MnSO₄ for 30 min. *p* values were determined using one-way ANOVA with Dunnett’s correction for multiple comparisons against the MOPS control. Error bars represent mean ± SD; *n* = 3. Each autoradiogram is representative of at least three independent experiments.

**Fig. S2.**
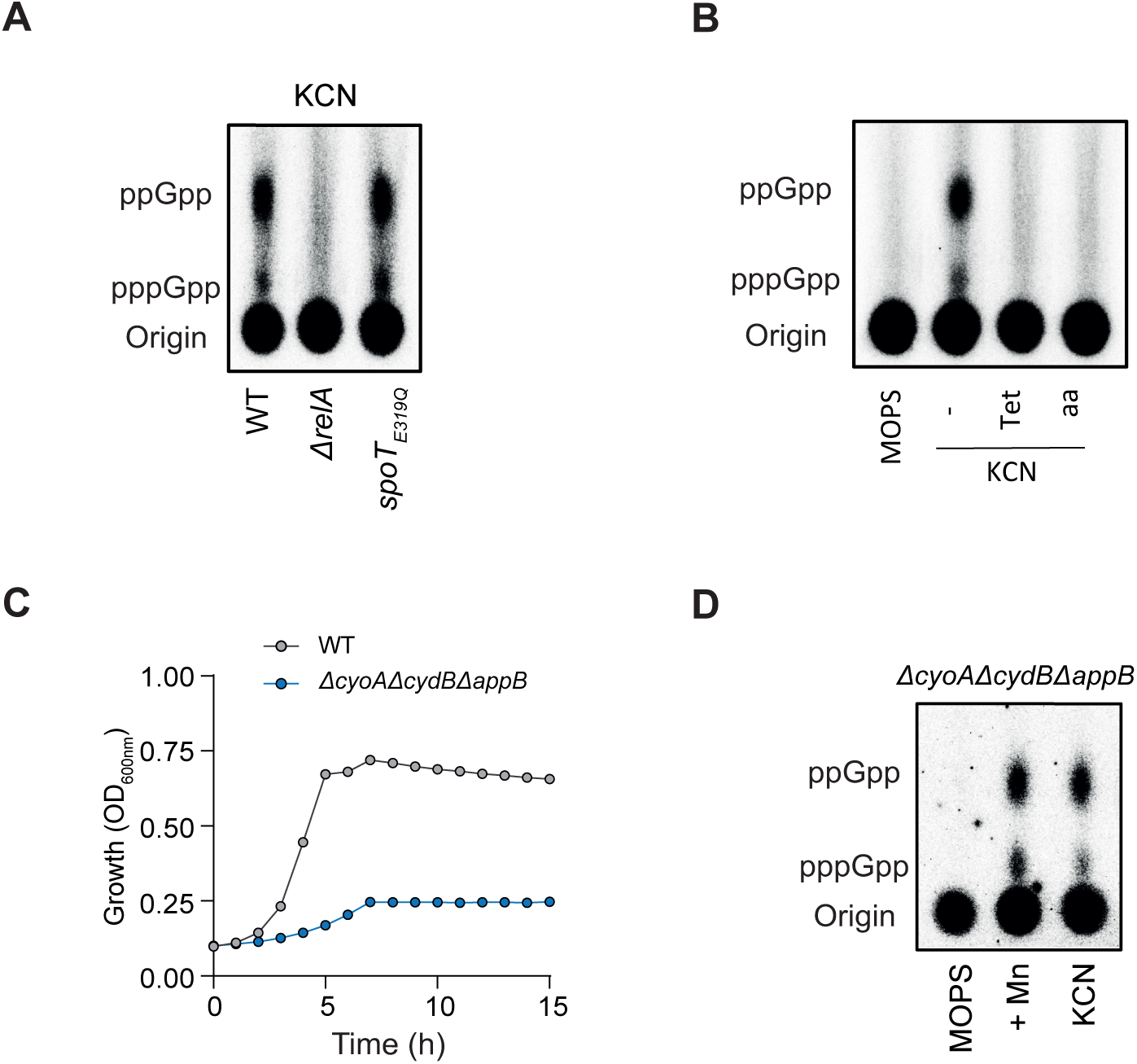
Mn and KCN treatment lead to (p)ppGpp accumulation independently of cytochrome oxidase. **(A)** Representative TLC autoradiogram showing nucleotide content in wild-type (WT), Δ*relA*, and *spoT*_E319Q_ strains after treatment with 1 mM KCN for 30 min. **(B)** Representative TLC autoradiogram of nucleotide content in WT cells incubated for 30 min in MOPS minimal medium (unstressed) or treated with 1 mM KCN, with or without additional supplementation: no treatment (−), tetracycline (Tet), or a mixture of 20 amino acids (aa). **(C)** Growth curves of the WT and the Δ*cyoA*Δ*cydB*Δ*appB* strains in MOPS minimal medium. **(D)** Representative TLC autoradiogram of nucleotide content in Δ*cyoA*Δ*cydB*Δ*appB* strain grown in MOPS minimal medium with or without 1 mM MnSO₄ (+Mn) or 1 mM KCN for 30 min.

**Fig. S3.**
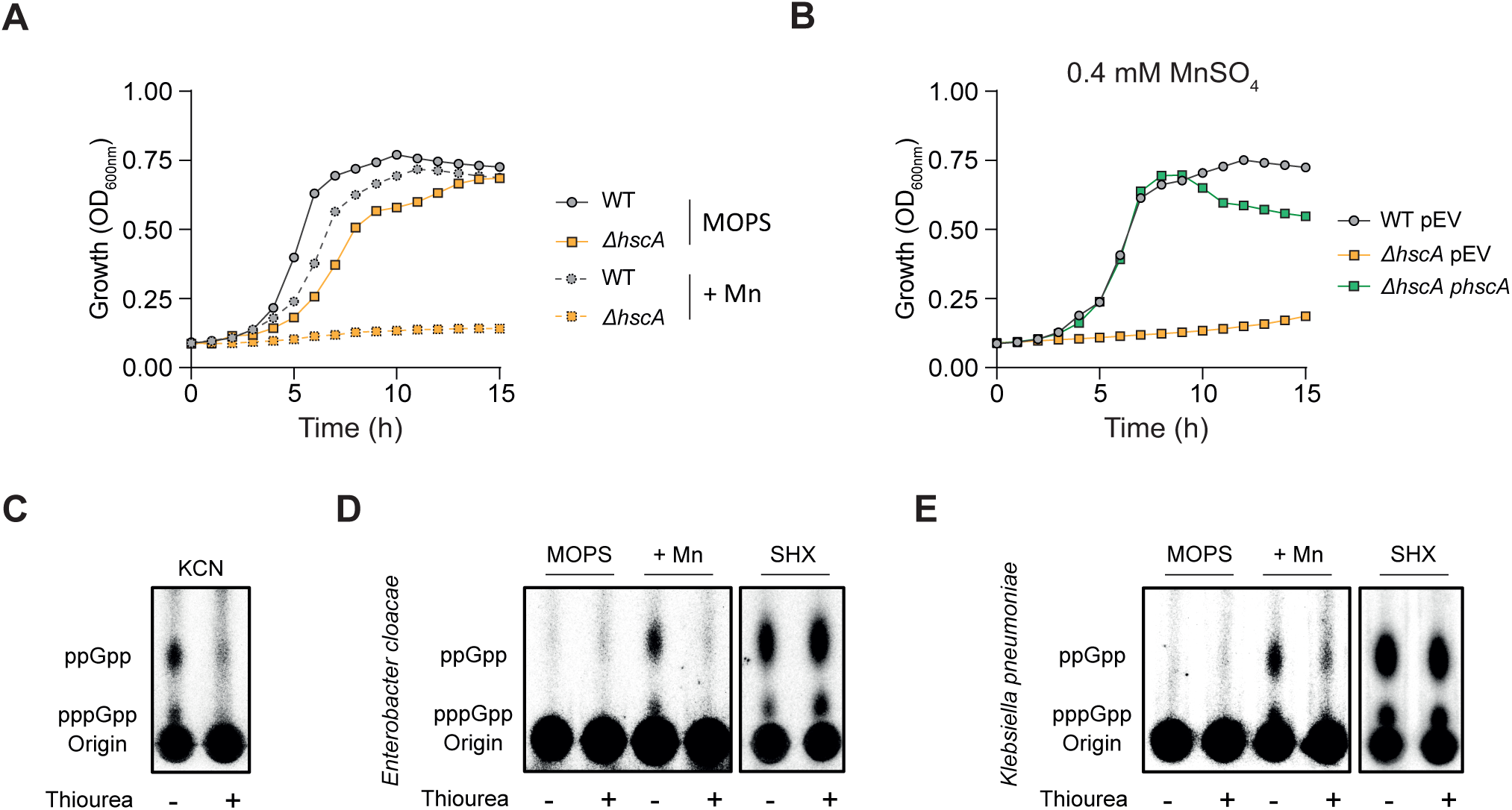
Manganese intoxication leads to Fe-S cluster damage. **(A)** Growth curves of the WT and the Δ*hscA* strains in MOPS minimal medium in presence or absence of 0.4 mM MnSO_4_ (+Mn). **(B)** Growth curves of the WT or Δ*hscA* strain complemented with an empty vector (pEV) or with *hscA* (p*hscA*) in MOPS minimal medium supplemented with 0.4 mM MnSO₄. **(C)** Representative TLC autoradiograms of ³²P-labeled nucleotide extracts from WT *Salmonella* grown for 30 min in MOPS minimal medium in presence of 1 mM KCN and supplemented with (+) or without (-) 100 mM thiourea. **(D-E)** Representative TLC autoradiograms of ³²P-labeled nucleotide extracts from WT *Enterobacter cloacae* (**D**) and *Klebsiella pneumoniae* (**E**) grown for 30 min in MOPS minimal medium in presence or absence of 1 mM MnSO₄ (+Mn) or 0.4 mg/mL SHX and supplemented with (+) or without (-) 100 mM thiourea.

**Fig. S4.**
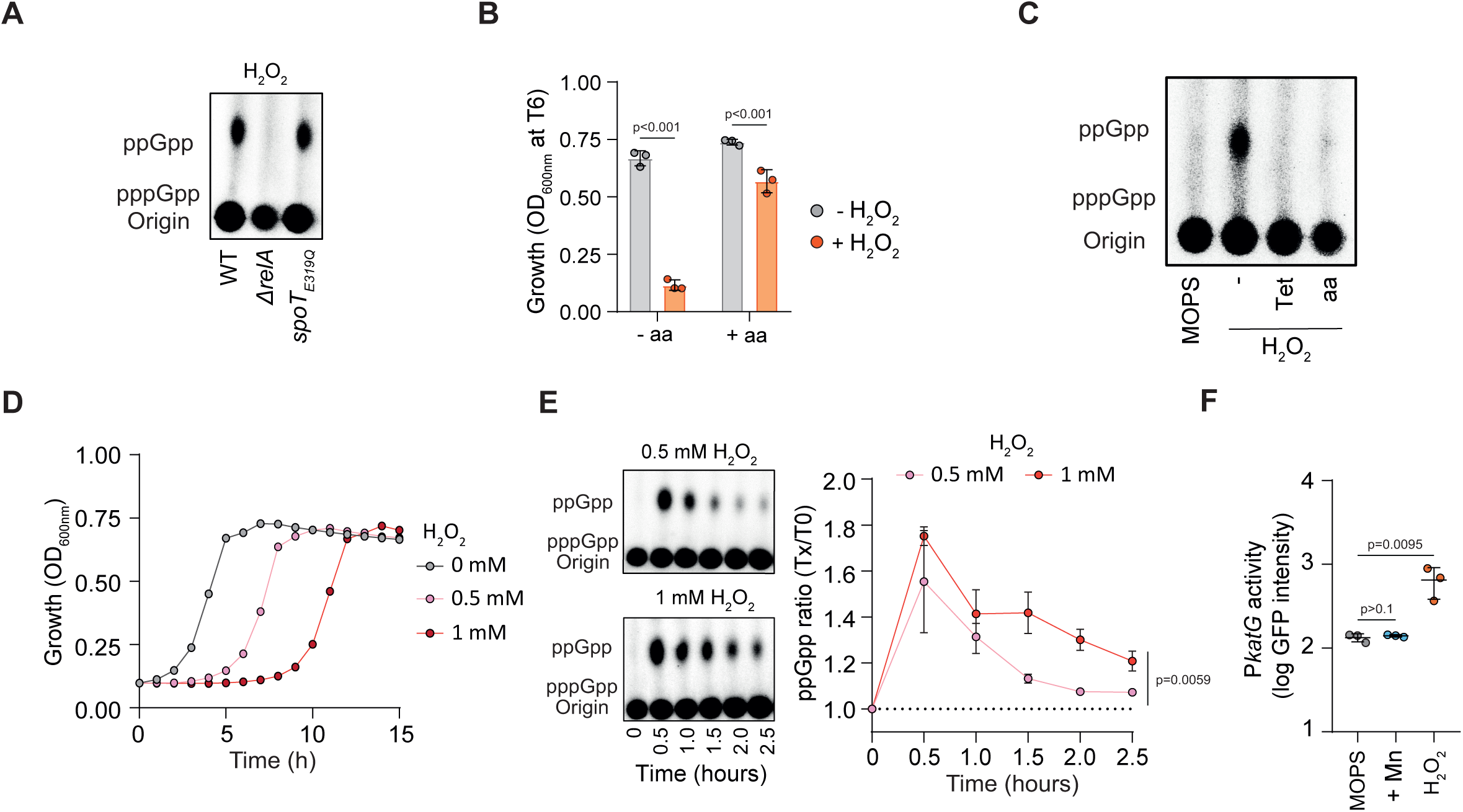
Oxidative stress induces (p)ppGpp via RelA activation in response to amino acid limitation. **(A)** Representative TLC autoradiogram showing nucleotide content in WT, Δ*relA*, and *spoT*_E319Q_ strains after treatment with 1 mM H₂O₂ for 30 min. **(B)** Optical density (OD₆₀₀_nm_) of WT *Salmonella* grown in MOPS minimal medium with or without 1 mM H₂O₂ and supplemented with (+aa) or without (−aa) a mixture of all 20 amino acids. **(C)** Representative TLC autoradiogram of nucleotide content in WT cells incubated for 30 min in MOPS minimal medium (unstressed) or treated with 1 mM H₂O₂, with or without additional supplementation: no treatment (−), tetracycline (Tet), or a mixture of 20 amino acids (aa). **(D)** Growth curves of WT *Salmonella* in MOPS minimal medium supplemented with 0, 0.5, or 1 mM H₂O₂. **(E)** Representative TLC autoradiograms (left) and quantification (right) of ³²P-labeled nucleotide extracts from WT *Salmonella* over time following addition of 0.5 or 1 mM H₂O₂. **(F)** Quantification of *PkatG_gfpOVA* reporter activity in MOPS minimal medium in absence (MOPS) or presence of 1 mM MnSO₄ (+Mn) or 1 mM H₂O₂ for 30 min. *p*-values were calculated using multiple *t*-tests (B) or using one-way ANOVA with Dunnett’s correction for multiple comparisons against the MOPS control (F). For (E), *p*-values were determined using an unpaired *t*-test at the final time point. Error bars represent mean ± SD; *n* = 3. Each autoradiogram is representative of at least three independent experiments.

**Fig. S5.**
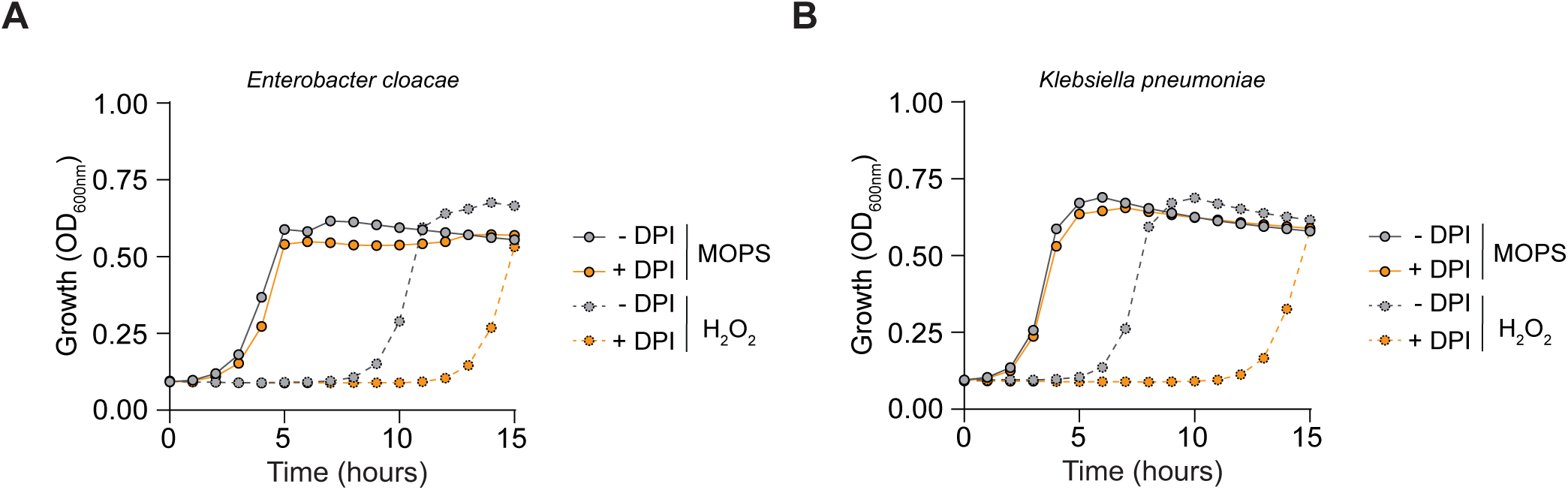
Iron availability modulates the lag phase of clinically relevant pathogens during oxidative stress. (A–B) Growth curves of WT *Enterobacter cloacae* (**A**) and *Klebsiella pneumoniae* (**B**) in MOPS minimal medium containing 0 or 0.5 mM H₂O₂, with or without 0.1 mM DPI.

**Fig. S6.**
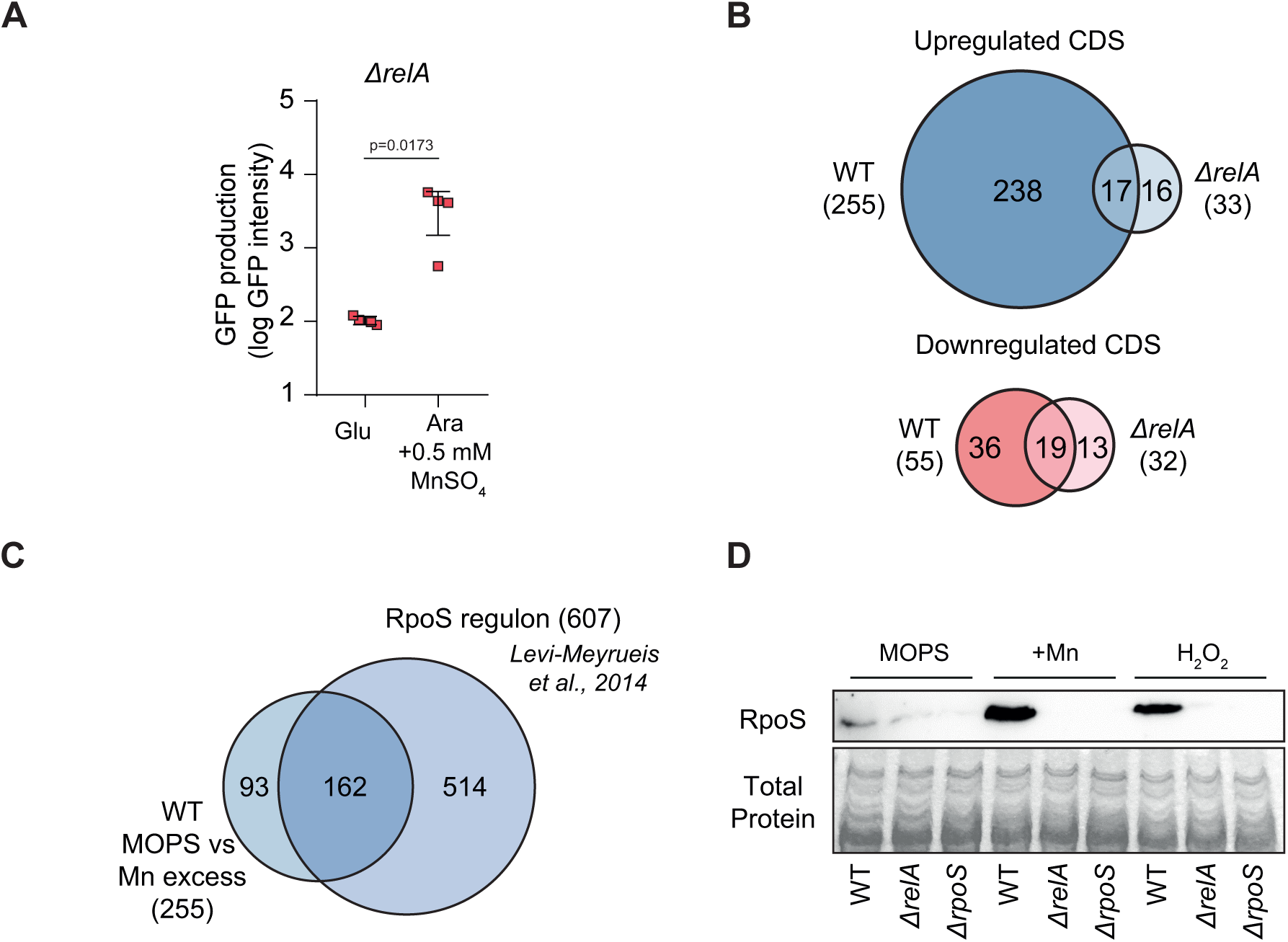
(p)ppGpp produced by RelA reshapes bacterial transcription in response upon Fe-S cluster damage. **(A)** Quantification of transcriptional and translational activity in *ΔrelA* strain grown in MOPS minimal medium with glucose (Glu) or arabinose (Ara), in the absence (MOPS) or presence (+Mn) of 0.5 mM MnSO₄. *p* values were determined using an unpaired *t*-test. Error bars represent mean ±SD; *n* ≥ 3. **(B)** Venn diagram showing the overlap between genes regulated in WT and *ΔrelA* strains during Mn stress. Numbers indicate the number of genes in each subset. **(C)** Venn diagram showing overlap between RpoS-regulated genes identified in (45) and those upregulated in this study. Numbers indicate the number of genes in each subset. **(D)** Immunoblot analysis of protein extracts from WT, Δ*relA*, and Δ*rpoS* strains grown in MOPS minimal medium with or without 1 mM MnSO₄ (+Mn) or 1 mM H₂O₂. Total protein was visualized by Coomassie blue staining as loading control.

**Fig. S7.**
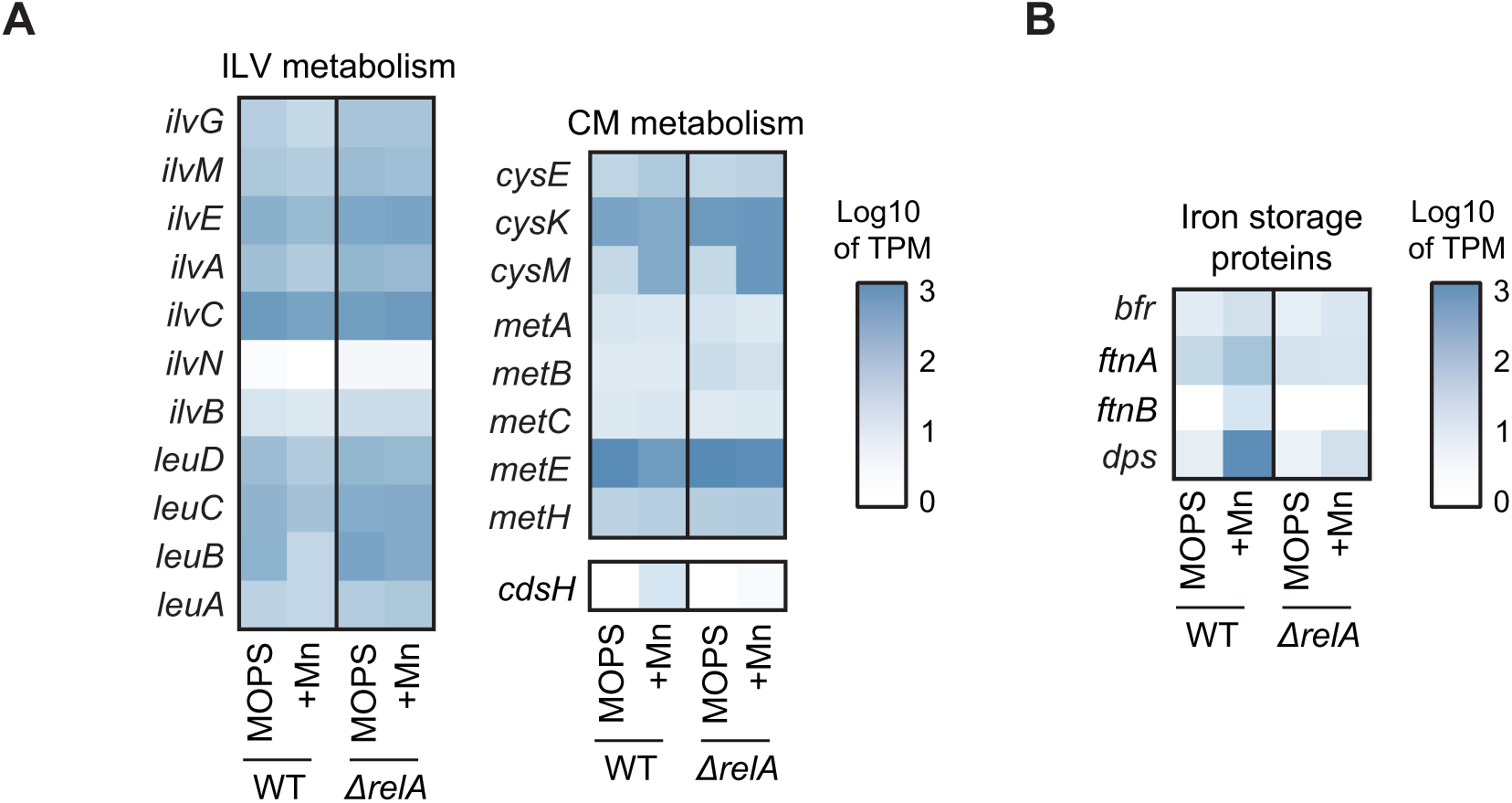
Impact of (p)ppGpp on gene expression involved in amino acid metabolism and iron homeostasis. **(A)** Heatmap showing expression of genes involved in ILV (branched-chain amino acid) and CM (sulfur-containing amino acid) metabolism in WT and *ΔrelA* strains in the absence (MOPS) or presence (+Mn) of 0.5 mM MnSO₄. **(B)** Heatmap showing expression of genes involved in iron storage in WT and *ΔrelA* strains in the absence (MOPS) or presence (+Mn) of 0.5 mM MnSO₄.

## Notes

### Competing Interest Statement

The authors have declared no competing interest.

