## Supplementary material for "Stress-Induced Iron-Sulfur Cluster Damage as a Conserved Trigger of the Stringent Response": Material & Methods

### MATERIAL AND METHODS

#### Bacterial strains and plasmids

Oligonucleotides, strains, and plasmids used in this study are listed in Supplementary Tables 1, 2, and 3. Additional construction details are provided in the Supplementary Methods. *E. coli* EC100D pir-116 was used for cloning purposes with pTOX2 and grown aerobically at 37°C in Luria–Bertani (LB) broth (Sigma) supplemented with 2% glucose. *Salmonella* strains were grown aerobically at 37°C in either LB or MOPS minimal medium (40 mM MOPS buffer, 0.4 mM tricine, 0.4 mM $K_2HPO_4$, 10 µM $FeSO_4$•$7H_2O$, 9.5 mM $NH_4Cl$, 276 µM $K_2SO_4$, 500 nM $CaCl_2$, 525 µM $MgCl_2$, 50 mM NaCl, 2.9 nM $(NH_4)_6Mo_7O_{24}$•$4H_2O$, 400 nM $H_3BO_3$, 30 nM $CoCl_2$, 9.6 nM $CuSO_4$, 80.8 nM $MnCl_2$, and 9.74 nM $ZnSO_4$; pH 7.2), supplemented with 0.2% glucose as the sole carbon source (referred to as MOPS throughout the manuscript). When indicated, the medium was supplemented with additional compounds at the following concentrations: amino acid (aa) mixture (40 µg/ml of each aa), thiourea (100 mM), reduced glutathione (50 mM), ascorbic acid (50 mM), serine hydroxamate (SHX, 0.4 mg/ml), $MnSO_4$ (0.5 or 1 mM), $H_2O_2$ (0.5, 1, or 2 mM), $Mg^{2+}$ (0.1 or 1 mM), $Fe^{2+}$ (0.1 or 1 mM), $Zn^{2+}$ (0.1 or 1 mM), $Cu^{2+}$ (0.1 or 1 mM), $Co^{2+}$ (0.1 or 1 mM), $MnCl_2$ (1 mM), N,N,N=,N=-tetrakis(2-pyridylmethyl)ethylenediamine (TPEN, 200µM), bathocuproinedisulfonic acid disodium salt (BCS, 100 mM) and 2,2'-dipyridyl (DPI, 0.1 mM or 1 mM). Electrocompetent cells were used for the transformation of *E. coli* and *Salmonella*. Antibiotics were used at the following concentrations where appropriate: ampicillin (100 µg/ml), kanamycin (50 µg/ml), and chloramphenicol (34 µg/ml).

Genomic mutations were construct with pTOX2 vectors as previously described (1). Briefly, the *E. coli* donor strain was mixed with the *Salmonella* recipient strain at a 1:5 or 1:20 ratio in a total volume of 1 ml. Cells were centrifuged at 8000 rpm for 2 minutes, and the supernatant was discarded. The remaining 100 µl was plated on LB agar containing 0.3 mM diaminopimelic acid (DAP) and 2% glucose and incubated at 37°C for 4 hours. Subsequently, cells were scraped and restreaked on LB plates containing 2% glucose and chloramphenicol. Single *Salmonella* colonies were then restreaked on LB with 2% rhamnose. Finally, chloramphenicol-sensitive ($Cam^S$) clones were screened by PCR.

**Growth assays**

Bacteria were grown overnight in MOPS. The overnight culture (10 µl) was diluted into 500 µl of fresh MOPS, supplemented with additional compounds when indicated. 200 µl of each sample was transferred into a flat-bottom 96-well plate (Greiner). Optical density at 600 nm ($OD_{600}$) was measured every hour for 15 hours at 37°C using a plate reader (BioTek Epoch 2). For panel 4A, *Salmonella* strains (WT or *ΔrelA*) were grown overnight in MOPS. Then, 10 µl of the overnight culture were diluted in 500 µl of MOPS supplemented with $MnSO_4$ (0, 0.5 or 1 mM) or $H_2O_2$ (0, 1 or 2 mM) in 1.5 ml eppendorf. Samples were incubated in thermal shaker at 37°C with shaking (1000 rpm) for 15 hours before $OD_{600}$ measurement.

**Measurement of (p)ppGpp**

Bacteria were grown overnight in MOPS. The overnight culture was then diluted 1:50 in fresh MOPS (100 µl of overnight culture into 5 ml) and incubated aerobically at 37°C for 90 min. Next, 20 µl of a 1 mCi $KH_2{}^{32}PO_4$ solution was added to 480 µl of bacterial culture, and the cells were incubated aerobically at 37°C for 60 min. Following incubation, cells were pelleted by centrifugation at 13,000 rpm for 3 min and resuspended in 25 µl of MOPS. For stress treatment, 5 µl of resuspended cells were diluted in 100 µl of MOPS containing the indicated stress conditions and incubated at 37°C with shaking for 30 min. After incubation, 40 µl of ice-cold 50% formic acid was added to each sample, thoroughly mixed, followed by an additional 30-minute incubation on ice. Samples were then centrifuged at 13,000 rpm for 5 minutes and then returned to ice. For thin-layer chromatography (TLC), 2.5 µl x4 of each sample was spotted onto a polyethyleneimine (PEI) cellulose plate (Macherey-Nagel). The PEI plates were developed in 1.25 M $KH_2PO_4$ (pH 3.4) at room temperature. Finally, TLC plates were air dried, wrapped in plastic and exposed against MS Storage Phosphor Screen (GE Healthcare) overnight. Phosphor screen were visualized using an Amersham Typhoon (Cytiva). Spot intensities were quantified with ImageJ software. Background was always subtracted from the original value before further calculation.

**Fluorescence accumulation in *Salmonella***

*Salmonella* strains (WT or Δ*relA*) containing the pFCcGi reporter were grown overnight in MOPS with ampicillin. Bacteria were then diluted (1:50) in MOPS containing 0.2 %

glycerol instead of glucose and grow aerobically for 90 min at 37°C. Then, media was supplemented with $MnSO_4$ (0.5 or 1 mM) or $H_2O_2$ (0.5 or 1 mM). After 30m of stress, 0.2 % arabinose was added to the media to induce the production of GFP. Each sample was collected after 30 min of induction and stored at 4°C in PBS prior analysis on an Attune NxT flow cytometer.

**Promoter activity assay**

*Salmonella* strain containing the P*katG_gfpOVA* was grown overnight in MOPS and ampicillin. Bacteria were then diluted in MOPS (20 µl overnight in 1 ml) and grow aerobically for 90 min at 37°C. Then, media was supplemented with 0.5 mM $H_2O_2$ and/or 0.1 mM DPI. Each sample was collected after 30 min of stress exposure and stored at 4°C in PBS prior analysis on an Attune NxT flow cytometer.

**RNA-Seq and analysis**

*Salmonella* strains (WT or *ΔrelA*) were grown overnight in MOPS in three biological replicates. Bacteria were then diluted in MOPS (1:50) and grow aerobically for 90 min at 37°C to reach an $OD_{600}$ of 0.2-0.3 in 50 ml. Then, media was supplemented with 0.5 $MnSO_4$. After 30 min of stress, bacteria were centrifuged and the pellets froze at -80°C. RNA extraction was performed using the zymo kit for quick RNA extraction. DNase treatment was carried out using the Turbo DNase kit from Invitrogen. After quality checking with a TapeStation (Agilent), RNA samples were sent to Genewiz for rRNA depletion, followed by cDNA library preparation and strand-specific RNA sequencing. Raw data were paired, trimmed, and mapped against the reference genome of *Salmonella enterica* serovar Typhimurium (accession numbers CP001362 and CP001363) using CLC software (Qiagen). Differential expression analysis (DE) and volcano plots were generated using CLC, with a chosen cutoff of log2 fold change and FDR < 0.05 to narrow down the deregulated transcripts. Unsupervised heatmaps were generated using the mean of Transcript Per Million (TPM) in $log_{10}$ of each condition and plotted using R package ggplot2(2).

RNA-seq analysis comparing MOPS medium to excess manganese (0.5 mM $MnSO_4$) is presented in **Table S4** (WT) and **Table S5** (Δ*relA*).

**Immunoblot analysis**

*Salmonella* strains (WT, Δ*relA*, and Δ*rpoS*) were grown overnight in MOPS medium. Crude protein extracts were prepared by inoculating fresh MOPS medium (1:50 dilution) and incubating for 90 minutes, followed by exposure to either 1 mM $MnSO_4$ or 1 mM $H_2O_2$ for 30 minutes. Cells were then resuspended in SDS–PAGE loading buffer and lysed by heating at 90 °C for 10 minutes. Proteins were separated by electrophoresis on a 12% SDS–polyacrylamide gel, transferred to a nitrocellulose membrane, and immunoblotted for 3 hours using a primary α-RpoS antibody (1:5,000; BioLegend Europe BV) and a secondary anti-mouse HRP-conjugated antibody (1:5,000; GE Healthcare). Detection was performed using Western Lightning Plus-ECL chemiluminescence reagent (Bio-Rad) and visualized with an ImageQuant LAS400 system (GE Healthcare).

#### Supplementary Tables

**Supplementary Table 1.** Oligonucleotides used in this study. Restriction sites are indicated in capital letter.

| Name | Sequence |
|---|---|
| **oSR904** | atggttctgttcgagcgcat |
| **oSR905** | gctaatgcggctttgctgaa |
| **oSR911** | tccccgggtaGCGGCCGCgcgctagtttcggttacggg |
| **oSR927** | accagaaccagaaccAAGCTTcttcggatcaaattcaccagc |
| **oSR928** | ggttctggttctggtAAGCTTccggatgtgattgatgcacg |
| **oSR929** | tccccgggtaGCTAGCaccagcatcctgattctggc |
| **oSR962** | ggacttgaacaggaacgccgtgcg |
| **oSR963** | cgttcctgttcaagtccagatccgtaccg |
| **oSR964** | tccccgggtaGCTAGCgtgattcgattgcgcaacgg |
| **oSR969** | aagtaactaaGAGCTCtactggaaacggtatctggc |
| **oSR976** | aagtaactaaGAGCTCcggtaaaagcgttgccgagc |
| **oSR999** | cccgcacggcgttcctgttc |
| **oSR1265** | ttaccgcagcgataaagcgg |

| | |
|---|---|
| **oSR1266** | cacaagcgtttcgcatgacg |
| **oSR1312** | cgataaactgcgaagcagcg |
| **oSR1313** | tcaatctcgatgccgttgcg |

**Supplementary Table 2.** Plasmids used in this study.

| Name | Description | Reference |
|---|---|---|
| **pRSc23** | pCA24N (pEV) | (3) |
| **pRSc30** | pCP20 | (4) |
| **pRSc35** | pFCcGi | (5) |
| **pRSc51** | pNB23 - P*katG_gfpOVA* | (6) |
| **pRSc139** | pTOX2_*ΔrelA* | This study |
| **pRSc145** | pTOX2_$spoT_{E319Q}$ | This study |
| **pRSc470** | pTOX2 | (1) |
| **pRSc531** | pCA24N_*hscA* (p*hscA*) | (3) |

**Supplementary Table 3.** Strains used in this study.

| Name | Description and relevant genotype | Source |
|---|---|---|
| **RSc1** | *Salmonella enterica* serovar Typhimurium strain 12023/14028 | ATCC 14028 |
| **RSc35** | *Salmonella enterica* serovar Typhimurium strain 12023/14028 pFCcGi | (7) |
| **RSc53** | *Salmonella enterica* serovar Typhimurium strain 12023/14028 *araAB::mcherry* P*katG_gfpOVA* | (7) |
| **RSc139** | MFD-pir pTOX2_*ΔrelA* | This study |
| **RSc145** | MFD-pir pTOX2_$spoT_{E319Q}$ | This study |

| | | |
|---|---|---|
| **RSc158** | *Salmonella enterica* serovar Typhimurium strain 12023/14028 Δ*relA* | This study |
| **RSc162** | *Salmonella enterica* serovar Typhimurium strain 12023/14028 $spoT_{E319Q}$ | This study |
| **RSc163** | *Salmonella enterica* serovar Typhimurium strain 12023/14028 Δ*relA* $spoT_{E319Q}$ | This study |
| **RSc329** | *Klebsiella pneumoniae subsp. pneumoniae* | ATCC 700603 |
| **RSc332** | *Enterobacter cloacae subsp. cloacae* | ATCC 13047 |
| **RSc364** | *Salmonella enterica* serovar Typhimurium strain 12023/14028 Δ*relA* pFCcGi | This study |
| **RSc393** | *Salmonella enterica* serovar Typhimurium strain 12023/14028 Δ*cyoA* ΔappB cydD::kan | (7) |
| **RSc465** | *Escherichia coli* EC100D pir-116 (reference strain for R6K plasmid propagation) | Epicenter Biotechnologies |
| **RSc466** | *Escherichia coli* MFD-pir (reference strain for conjugation) | (8) |
| **RSc515** | *Salmonella enterica* serovar Typhimurium strain 12023/14028 *rpoS::kan* (referred as Δ*rpoS*) | (9) |
| **RSc529** | *Salmonella enterica* serovar Typhimurium strain 12023/14028 Δ*hscA* | This study |
| **RSc530** | *Salmonella enterica* serovar Typhimurium strain 12023/14028 pCA24N | This study |
| **RSc531** | *Salmonella enterica* serovar Typhimurium strain 12023/14028 pCA24N_*hscA* | This study |
| **RSc532** | *Salmonella enterica* serovar Typhimurium strain 12023/14028 Δ*hscA* pCA24N | This study |
| **RSc533** | *Salmonella enterica* serovar Typhimurium strain 12023/14028 Δ*hscA* pCA24N_*hscA* | This study |

**Supplementary Table 4.** Differential gene expression in wild-type *Salmonella* grown in MOPS medium with or without 0.5 mM $MnSO_4$.

**Supplementary Table 5.** Differential gene expression in *ΔrelA Salmonella* grown in MOPS medium with or without 0.5 mM $MnSO_4$.

### Supplementary Methods

#### Description of Plasmids

**pRSc139** (pTOX2_*relA*)

Upstream and downstream regions of the *relA* gene from the wild-type *Salmonella enterica* serovar Typhimurium strain 12023/14028 were amplified using primers oSR969–927 and oSR928–929, and cloned into pTOX2 using the NheI and SacI restriction sites.

**pRSc145** (pTOX2_ $spoT_{E319Q}$)

DNA fragments of the *spoT* gene from strain 12023/14028 encompassing the E319Q mutation site were amplified using primers oSR976–962 and oSR963–964, and cloned into pTOX2 using the NheI and SacI restriction sites.

#### Description of Strains

**RSc158** (*12023/14028 ΔrelA*)

Biparental mating between strain *Salmonella enterica* serovar Typhimurium 12023/14028 (RSc1) and the diaminopimelic acid (DAP) auxotroph MFD-pir carrying pTOX2_*ΔrelA* (RSc139) was performed. Selection was carried out on LB agar supplemented with chloramphenicol and glucose, followed by cultivation in LB with rhamnose. Chloramphenicol-sensitive ($Cam^S$) colonies were screened by PCR using primers oSR904 and oSR905.

**RSc162** (*12023/14028* $spoT_{E319Q}$)

Biparental mating between strain 12023/14028 (RSc1) and the DAP auxotroph MFD-pir carrying pTOX2_$spoT_{E319Q}$ (RSc145) was performed. Selection was done on LB agar containing chloramphenicol and glucose, followed by cultivation in LB with rhamnose. $Cam^S$ colonies were screened by PCR using primers oSR911 and oSR999, and the presence of the E319Q mutation was confirmed by sequencing.

**RSc163** (*12023/14028 ΔrelA* $spoT_{E319Q}$)

Biparental mating between strain RSc158 (12023/14028 *ΔrelA*) and the DAP auxotroph MFD-pir carrying pTOX2_*spoT*$_{E319Q}$ (RSc145) was conducted. Selection was performed on LB agar supplemented with chloramphenicol and glucose, followed by growth in LB with rhamnose. Cam$^{S}$ colonies were screened by PCR using primers oSR911 and oSR999, and the mutation was validated by sequencing.

**RSc515** (12023/14028 *rpoS::kan)*

The *rpoS::kan* strain from the Single-Gene Deletion (SGD) collection (9) was validated by using primers oSR1265-1266.

**RSc533** (12023/14028 *ΔhscA)*

The *hscA::kan* strain from the Single-Gene Deletion (SGD) collection (9) was transformed with pCP20 to obtain the *ΔhscA* strain which was subsequently validated using primers oSR1312-1313.
